# The effects of Paternal Radiofrequency Radiation Exposure on Sperm and Testicular Integrity in F1-Male Swiss Albino Mice

**DOI:** 10.1101/2024.01.23.576880

**Authors:** Neelam Pandey, Sarbani Giri, GD Sharma

## Abstract

The escalating prevalence of infertility and reproductive malignancies in the youth demographic warrants thorough investigation. The escalating ubiquity of communication devices, such as mobile phones and Wi-Fi, exposes individuals to unprecedented levels of Radiofrequency Radiation (RFR), raising concerns about its potential association with reproductive issues and cancer. In this study, we endeavored to elucidate the impact of RFR exposure on the fertility of F0 male Swiss albino mice, as well as the consequential effects on sperm quality and testicular morphology in F1 male mice during adulthood. F0 male mice were subjected to RFR exposure for 3 hours per day, twice daily, over a span of 35 days, followed by mating with naïve, unexposed females, culminating in a comprehensive endpoint evaluation. The repercussions of RFR exposure included a diminution in sperm count, heightened incidence of sperm defects, augmented DNA damage in both sperm and bone marrow cells, diminished sperm viability, and aberrations in testicular histology. Subsequent mating of exposed male mice with naïve females revealed no discernible effects on pregnancy outcomes, as indicated by the fertility index, litter size, and copulation index.Nevertheless, F1 male offspring exhibited a noteworthy reduction in sperm count and depletion of the germinal epithelium within seminiferous tubules in the testes. These findings propose that RFR exposure to F0 males may precipitate diminished sperm count in subsequent F1 generations, implicating the potential role of epigenetic modulation through the male germline.

## 1. Introduction

Infertility, impacting approximately 15% of reproductive-age couples, manifests as male factor infertility in about 50% of cases. Male reproductive dysfunction can stem from various levels, including pre-testicular (hypothalamic or pituitary damage), testicular (testis failure), post-testicular (sperm delivery obstruction), or a combination thereof. The etiology of male infertility, though multifactorial, often remains largely idiopathic (Hamada and Agarwal, 2012).

Radiofrequency (RF) radiation exposure has been linked to numerous effects, with a well-characterized impact primarily studied in exposed individuals or transplacental effects on fetuses of exposed mothers (Gul, 2009; Shahin, 2017). Notably, there is a dearth of intergenerational evidence regarding the paternal preconception exposure to RF radiations through the germline. The potential repercussions of paternal exposure to RF radiations are predominantly overlooked, highlighting a gap in our current understanding of the intergenerational impacts of such exposures.

Prior investigations, including studies by Gulati et al. (2018), have explored the impact of RF radiation on male reproductive functions in both humans and laboratory rodents. The consequences of inherited germline mutations can be profound for affected offspring and may carry significant socio-economic ramifications at the population level. Extensive evidence from animal studies establishes that environmental exposures have the potential to elevate mutation frequencies in germ cells (DeMarini, 2012). Notably, environmental factors such as RF radiation, identified as potential germ cell mutagens (Zalata, 2015), may contribute to the occurrence of sporadic genetic diseases in human populations by inducing de novo mutations. Presently, there is a limited understanding of lifestyle choices or environmental variables that lead to mutations in gametes transmitted to offspring or cause transgenerational genetic instability. This knowledge gap underscores the need for further research into the intricate relationship between environmental exposures, germline mutations, and their implications for the health of future generations.

The clinical implications of radiation-induced instability are currently constrained by a lack of understanding of the underlying mechanisms. The concept of ‘genetic memory,’ capable of destabilizing the non-exposed progeny of irradiated cells or organisms, may arise from a complex interplay of molecular, biochemical, and cellular events. Each of these events could potentially represent unknown pathway(s) in cellular stress response. While many pathways involved in the mammalian cellular response to radiation, such as DNA damage recognition, cell cycle arrest, and apoptosis, have been characterized (Pandey and Giri, 2018, 2017), there is evidence suggesting that the ability of cells to exhibit elevated mutation rates may not solely be attributed to conventional mechanisms but is likely associated with epigenetic events (Gul, 2009; Kumar, 2021). In this context, the present study unravels data on transgenerational changes in the germinal epithelium within seminiferous tubules in the non-exposed offspring of irradiated male mice. These findings offer valuable insights into the mechanisms of RF radiation-induced transgenerational changes associated with paternal exposure, shedding light on the intricate processes involved in cellular responses to radiation and their potential impact across generations.

## 2. Results

### 2.1. Sperm Defects in RFR exposed male mice

Scanning Electron Microscopy (SEM) analysis of spermatozoa disclosed distinctive morphological aberrations in mice subjected to Radiofrequency Radiation (RFR) exposure. Notable findings included the presence of blebs, notch-like protrusions, and specific intrusions in the plasma membrane. Additionally, the SEM images revealed the occurrence of short and stubby sperm with poorly formed surface structures in the exposed group (Fig 1B-1I). In contrast, these morphological defects were notably absent in the sperm of control mice, underscoring the specificity of the observed abnormalities in response to RFR exposure.

**Fig. 1.**
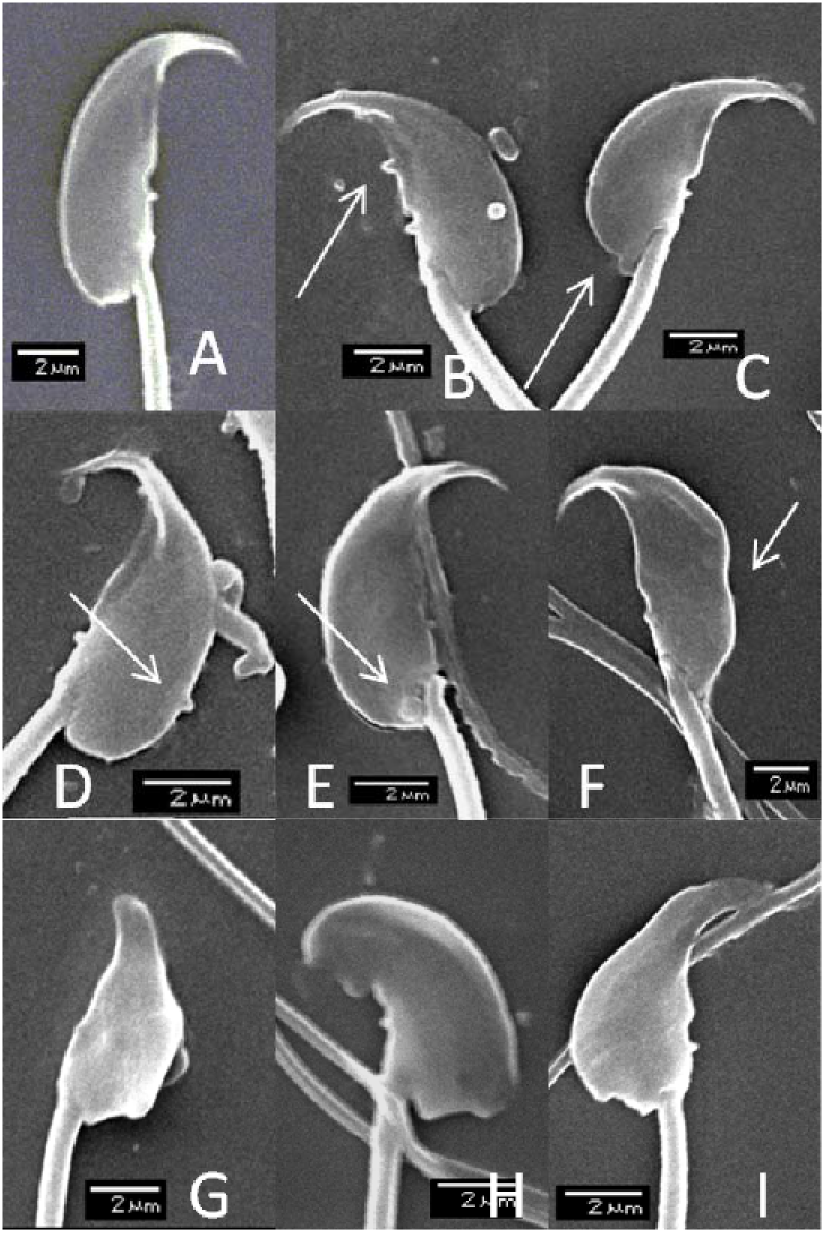
Scanning electron microscopy of epididymal sperms of normal and mice exposed to RFR for one spermatogenic cycle. In **panel A-**is the typical normal sperm from control mice; **panel B-D** – are seen sperms with short protruding structure (notch like) on borders (arrows); **panel E**-sperm with abnormal bleb at the neck region (arrow); **panel F**-abnormal sperm with intrusion on the membrane; **panel G-I**, are seen examples of aberrant head morphology i.e. short and stubby with poorly formed surface structures.

### 2.2 DNA damage in spermatozoa in RFR exposed male mice

Alkaline comet assay was performed to study the DNA integrity in sperm cells of F0 male mice after 35 days of exposure to RFR for 6 hr/day daily (Fig.2). Significant decrease in the sperm head DNA (%) with simultaneous increase in Tail DNA (%) and OTM was observed. Approximately, three time increase in damage index (DI) was seen in sperm cells from F0 male mice exposed to RFR for 35 days in comparison to control F0 mice. Damage Frequency (DI) represents the number of cells in each comet class. Control F0 male mice showed more than 60 percent cells in low DNA damage classes such as class0, 1 and 2 however, RFR exposed male showed more than 80 percent cells in high DNA damage classes such as class 3 and 4. Further, more than three times increase was observed in the class 4 (high damage) frequency.

**Fig. 2.**
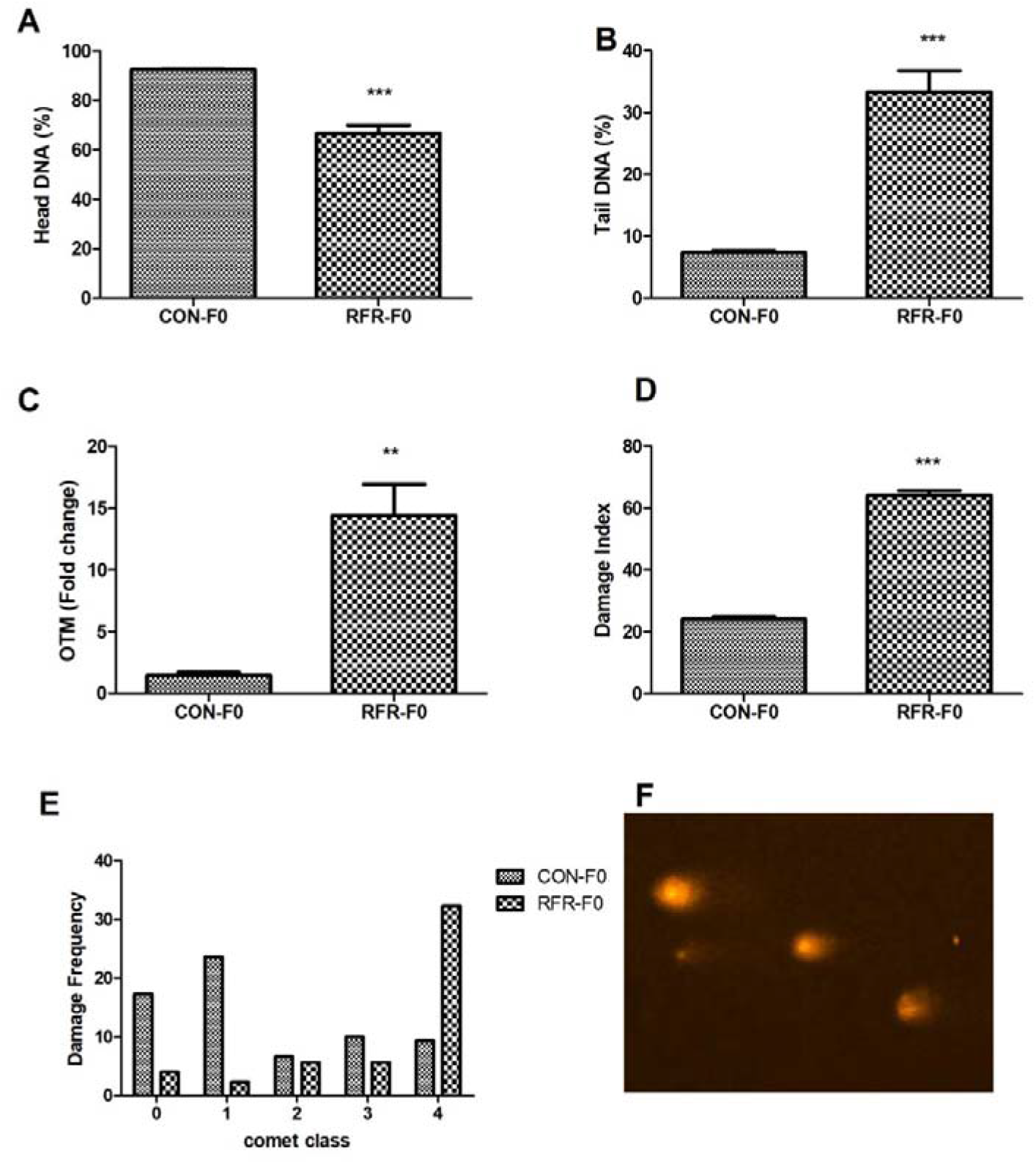
Comet assay in sperm cells of F0 male control and mice exposed to RFR for 35 days. (A) Percentage of Head DNA, (B) Percentage of Tail DNA, (C) fold change in Olive Tail Moment (OTM). (D) Damage Index (E) Damage frequency and (F) Gel picture showing EtBr stained comets in sperm cells. Significantly different from control at P < 0.001 (***), and P < 0.01(**). Each Bar represents the group mean ± SE (n=6).

### 2.3. DNA damage in bone marrow cells in RFR exposed male mice

Figure 3 provides a comprehensive summary of the comet assay outcomes conducted on bone marrow cells in F0 male mice following a 35-day exposure to Radiofrequency Radiation (RFR) for 6 hours per day. The results reveal a significant decrease in sperm head DNA percentage, accompanied by a simultaneous increase in Tail DNA percentage and Olive Tail Moment (OTM). Specifically, there was an approximately 2.5-fold increase in the Damage Index (DI) observed in sperm cells from F0 male mice exposed to RFR for 35 days compared to the control group. The Damage Frequency (DF) metric, representing the number of cells in each comet class, further elucidates the impact of RFR exposure. Control F0 male mice exhibited more than 60 percent of cells in low DNA damage classes such as class 0, 1, and 2. Conversely, RFR-exposed males displayed more than 80 percent of cells in high DNA damage classes, specifically class 3 and 4. Notably, there was a greater than threefold increase observed in the frequency of cells in class 4, indicative of high DNA damage. These findings underscore the substantial genotoxic effects induced by RFR exposure in the examined bone marrow cells.

**Fig. 3.**
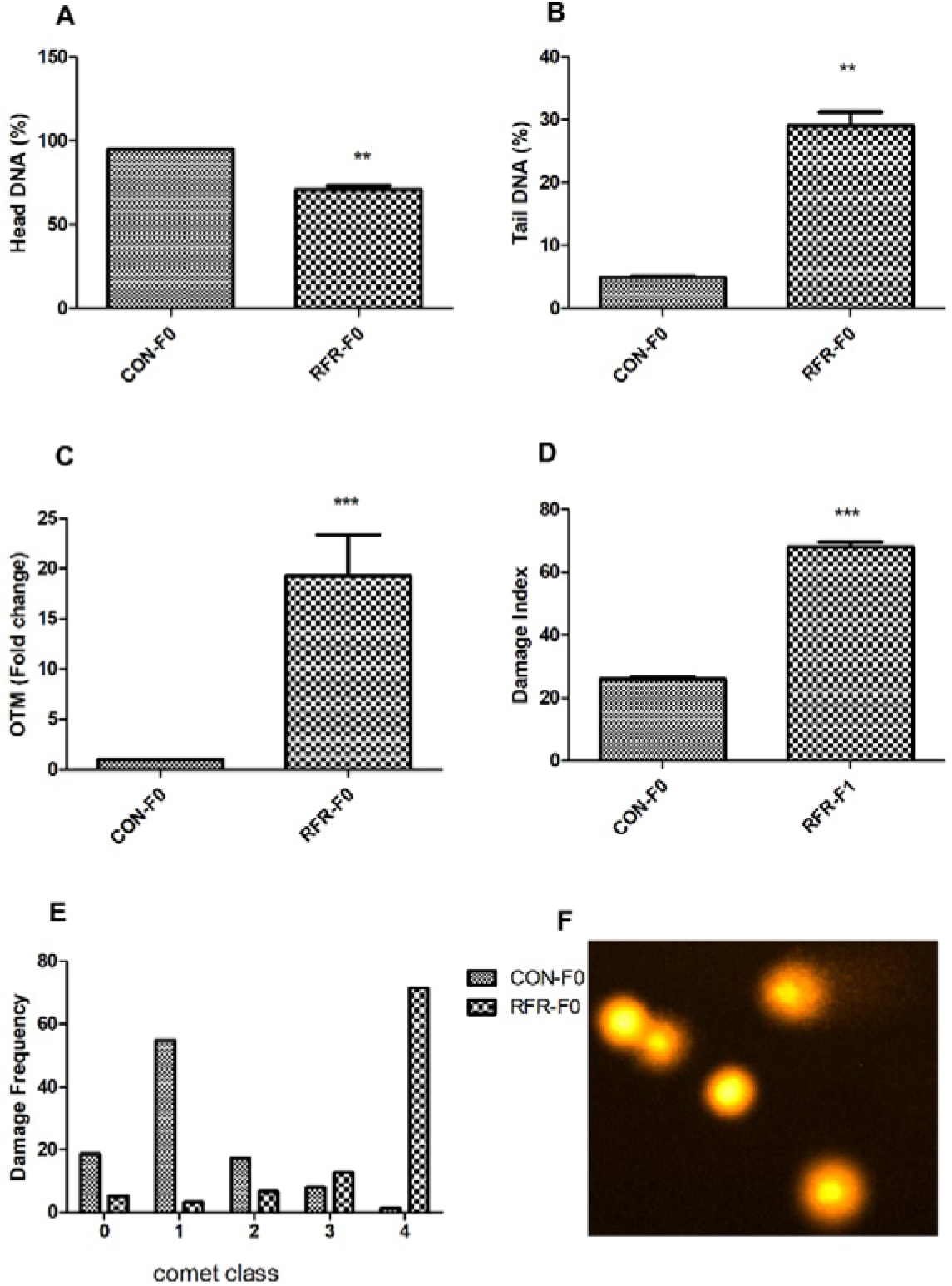
Comet assay in bone marrow cells of F0 male control and mice exposed to RFR for 35 days. (A) Percentage of Head DNA, (B) Percentage of Tail DNA, (C) fold change in Olive Tail Moment (OTM). (D) Damage Index (E) Damage frequency and (F) Gel picture showing EtBr stained comets in bone marrow cells. Significantly different from control at P < 0.001 (***), and P < 0.01(**). Each Bar represents the group mean ± SE (n=6).

### 2.4. Sperm viability in exposed F0 male mice

Live/dead sperm cell analysis was conducted in control, RFR-exposed F0, and RFR-exposed F1 male mice utilizing propidium iodide (PI) staining through flow cytometry (Fig. 4.A-B). The results indicate that F0 male mice exposed to RFR for 35 days exhibited a significant reduction in the percentage of live sperm compared to both the control group and RFR-exposed F1 male mice. Simultaneously, there was an increase in the percentage of dead sperm observed in RFR-exposed F0 mice. These findings suggest a notable impact of RFR exposure on sperm viability in F0 male mice, with potential implications for reproductive outcomes.

**Fig. 4.**
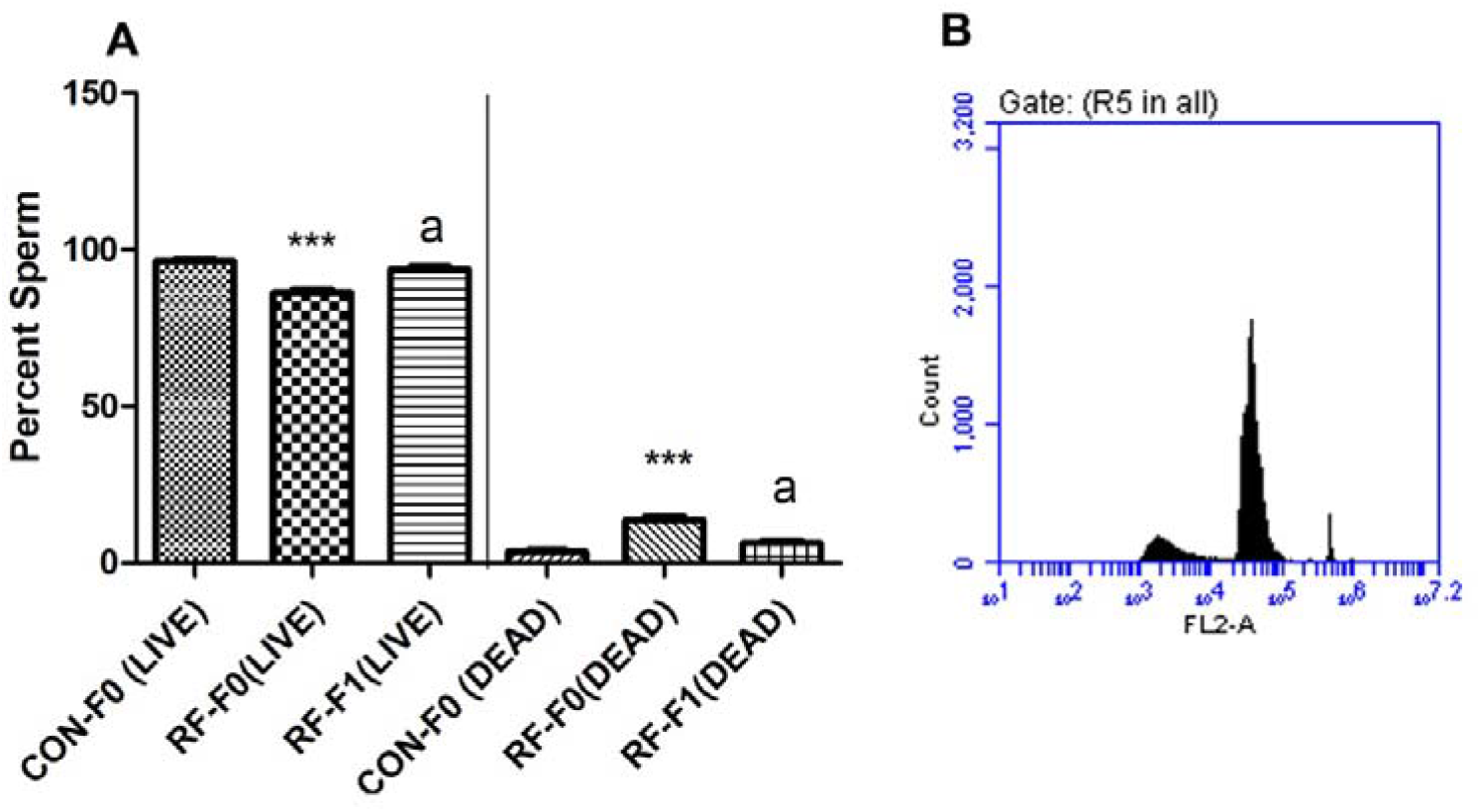
PI staining in sperm cells of F0 control, F0 and F1 male mice exposed to RFR for 35 days. (A) Percentage of live and dead cells, and (B) Representative cytogram showing peaks of live and dead sperms.

### 2.5. Mating studies in exposed F0 male mice

Control and RFR-exposed F0 male mice were paired with naive, unexposed females (Table 1). Notably, there was no statistically significant decrease observed in copulation index, fertility index, live birth index, or litter size. However, a significant increase in the male-to-female ratio was noted in the RFR-exposed F1 group compared to the F1 control group. Although no statistically significant difference was observed on Postnatal Day 1 (PND1) in the F1 generation survival index, a significant decrease in the survival of F1 pups was evident on PND 4, PND 7, PND 14, and PND 21 (Table 1). These findings suggest a potential impact of paternal RFR exposure on the male-to-female ratio and the survival of offspring during early postnatal development.

**Table 1.**
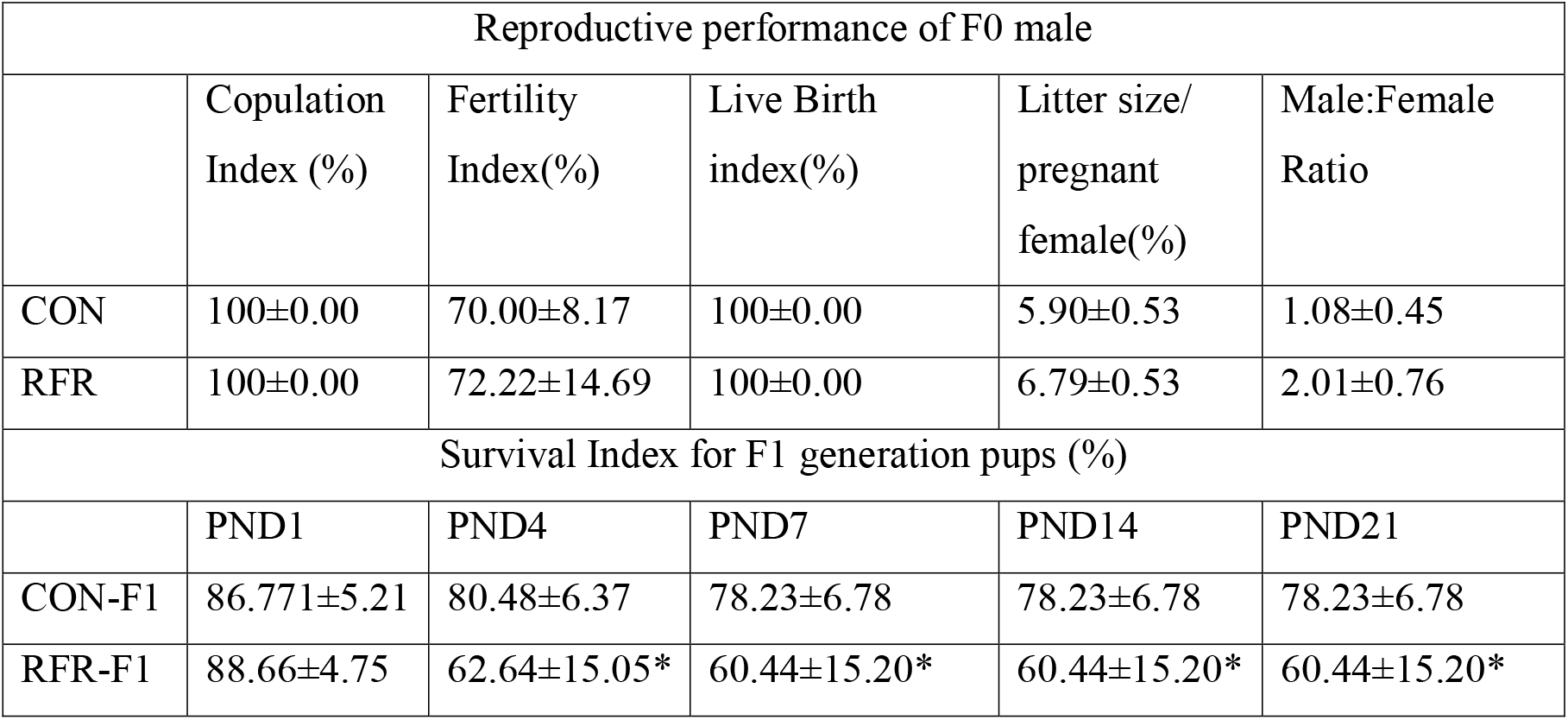
Effect of RFR exposure on reproductive performance of F0 male and survival of f1 progeny sired by unexposed female mated with control and RFR exposed F0 male.

### 2.6. Sperm count and sperm defects in F1 male at adulthood

Figure 5 illustrates the outcomes of reproductive organ weights, sperm count, and sperm head abnormality assays using a haemocytometer and eosin Y-stain, respectively. A noteworthy decrease was observed in the weights of primary reproductive organs, namely the testis and epididymis, in RFR-exposed F1 generation mice compared to both the control and RFR-exposed F0 mice (Fig 5A-B). There was a statistically significant increase, approximately 2.5 times, in sperm count in male mice exposed to RFR for 35 days compared to unexposed controls. Furthermore, a significant increase in the frequency of abnormal sperm heads was observed in F1 generation RFR-exposed mice compared to F1 generation control mice. However, it is noteworthy that the frequency of sperm head abnormalities decreased significantly, almost by half, in the F1 generation compared to F0 mice exposed to RFR for 35 days. Table 2 provides a summary of the types of abnormalities observed under a light microscope in RFR-exposed mice.

**Fig. 5.**
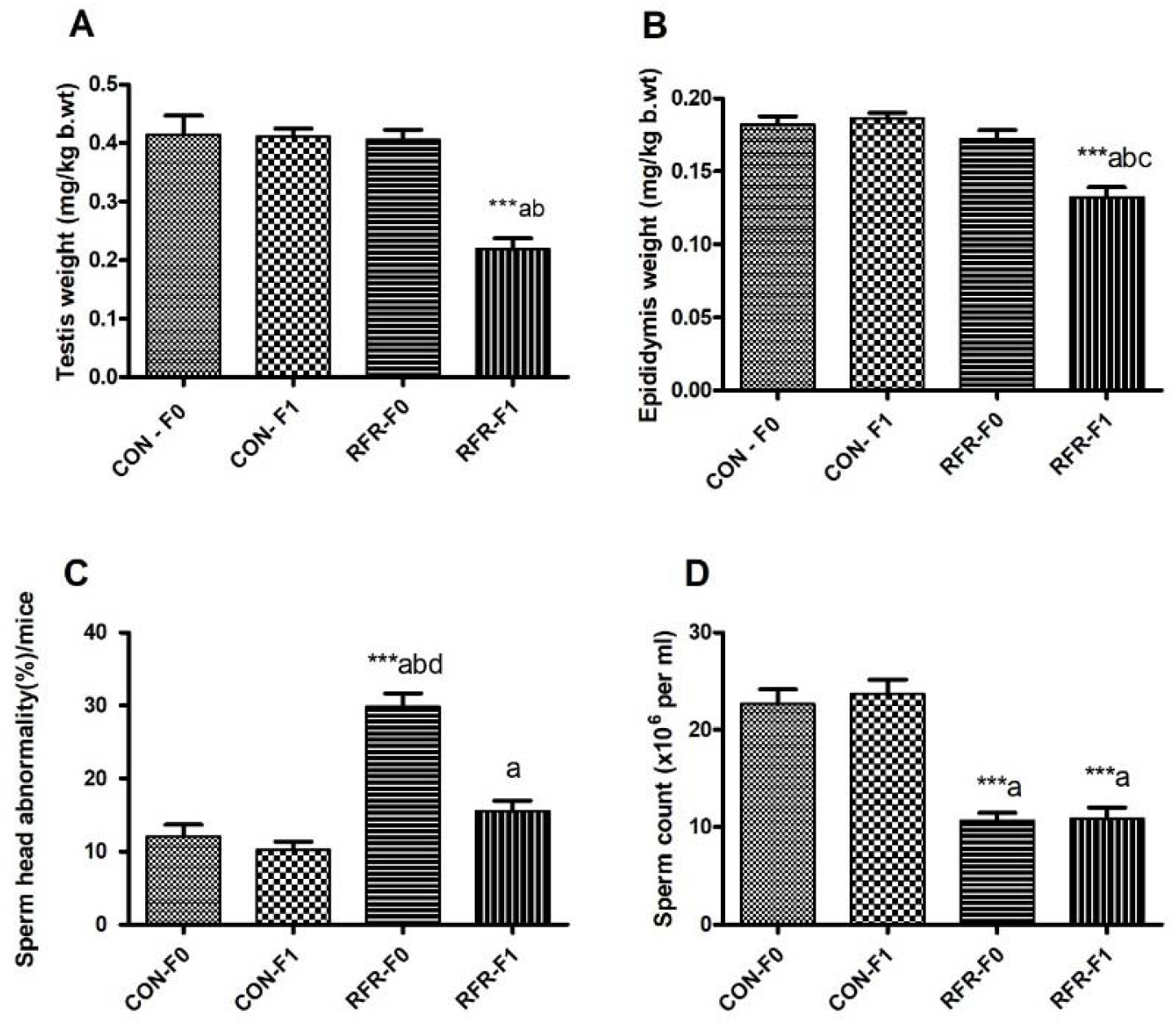
Figure showing relative weights of Testis (A), and Epididymal (B), Sperm head abnormalities(C) and Sperm count (million per ml) in F0 and F1 generation male in control and mice exposed to RFR for 35 days (n=6). Statistical test-one way ANOVA with tukey post hoc analysis. Each Bar represents the group mean ± SE (n=6). Asterisk indicates: *P ≤ 0.05, **P ≤0.01, ***P ≤ 0.001. ^a^ Control-F0 vs. control F1, RFR-F0, RFR-F1. ^b^ Control-F1 vs RFR-F0, RFR-F1. ^c^ RFR-F0 vs RFR-F1

**Table 2.**
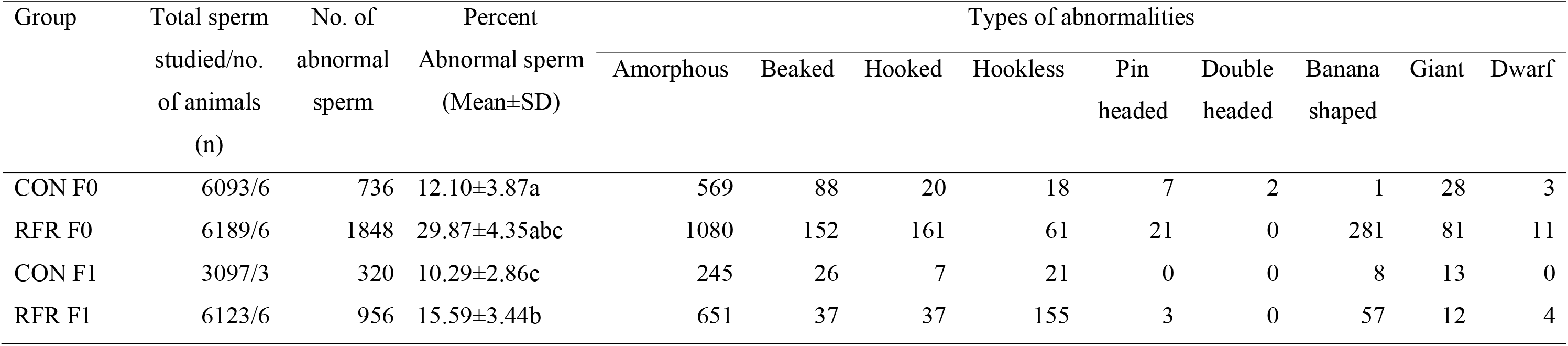
Incidence of sperm head abnormality in F0 and F1 generation control mice and mice exposed to RFR for 35 days and 35 days of post exposure recovery.

Additionally, a statistically significant decrease in sperm count was observed in both F0 and F1 generation mice exposed to RFR compared to control mice (Fig. 5D). These findings highlight the potential adverse effects of RFR exposure on reproductive parameters in subsequent generations.

### 2.7. Changes in Testis Histopathology of F1 male

Histological sections stained with Hematoxylin and Eosin (HE) staining revealed distinct changes, including an increased lumen size and intertubular space, demonstrating a thinning of the germinal epithelium (Fig. 6 C(a-b)). Morphometric analysis of the testis in both control and RFR-exposed mice in the F0 and F1 generations unveiled a significant decrease in the germinal epithelium area, coupled with a consistent increase in lumen size per seminiferous tubule per mouse in RFR-exposed mice compared to control (Fig. 6). Interestingly, RFR-exposed F1 generation mice also exhibited significantly altered germinal epithelium and lumen size compared to F1 control mice. These findings suggest that RFR exposure may induce structural changes in the testicular tissue, impacting both F0 and F1 generations.

**Fig. 6.**
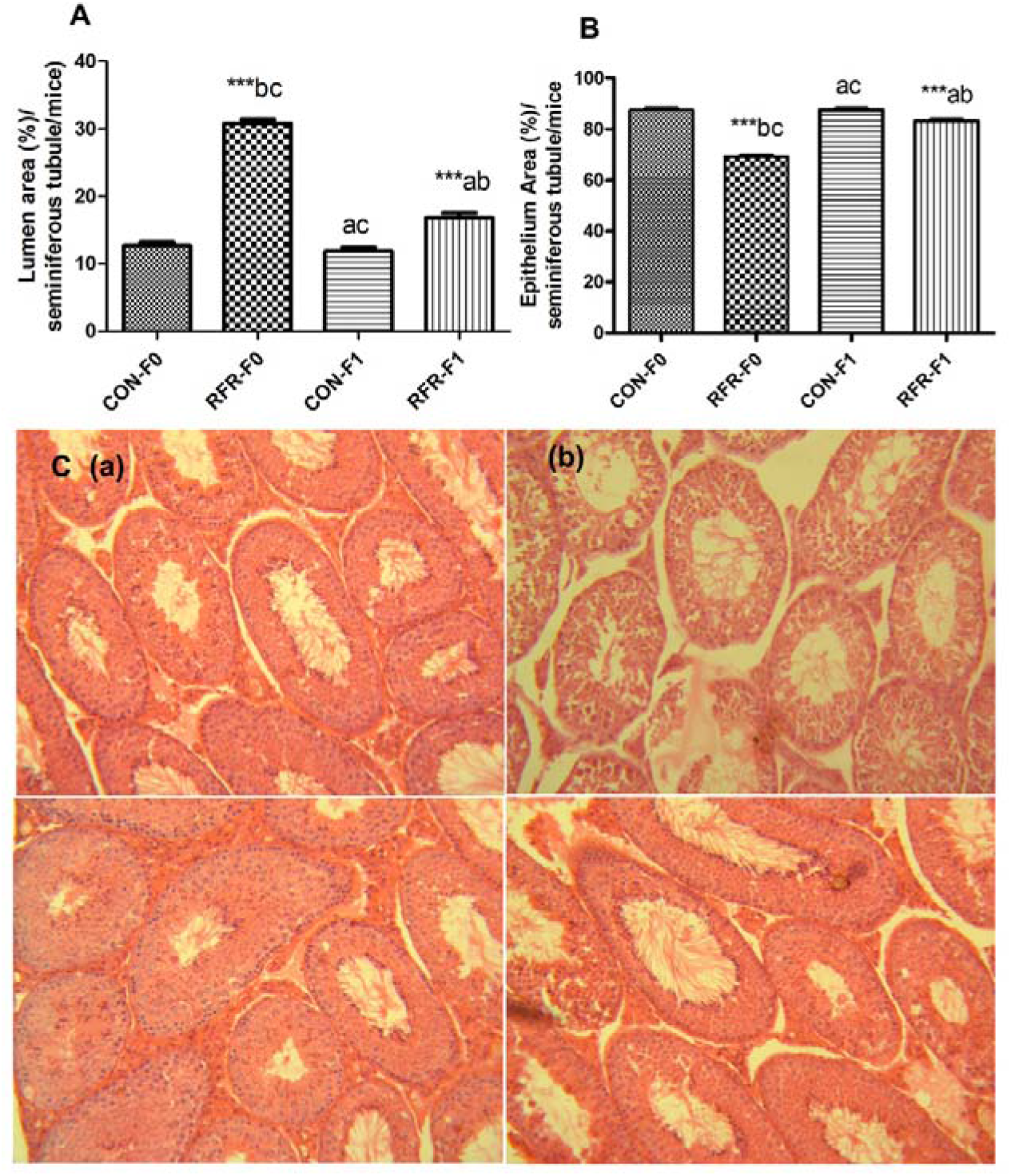
Exposure to RFR for 35 days altered testis histology of F0 and F1 male mice. (A) Percentage of lumen area and (B) seminiferous epithelium area per seminiferous tubule. C(a and c) seminiferous tubule exhibiting normal morphology with well arranged manner in testis of Control F0 and F1 generation male mice, C(b) increased lumen area and interstitial spaces between the seminiferous tubules in F0 male exposed to RFR for 35 days, and C (d) F1 generation male mice sired by unexposed female and exposed male.

## 3. Discussion

The present study explores the impact of sub-chronic exposure to Radiofrequency Radiation (RFR) on reproductive parameters in F0 mice and assesses the subsequent effects on sperm and testis characteristics in F1 progeny. Our findings reveal that whole-body exposure to RFR for 6 hours/day over 35 days induces alterations in testis histology, ultimately leading to a decrease in sperm count in the F1 generation. In line with our results, Gul et al. (2009) reported a reduction in the number of follicles in F1 female rats following RFR exposure to pregnant mothers. Additionally, RFR exposure is known to increase oxidative stress in mice placental tissue. Exposure to a 2.4 GHz frequency led to elevated lipid peroxidation, increased superoxide dismutase (SOD) activity, apoptosis, and overexpression of CDKN1A and GADD45a in mice placenta, potentially affecting placental gene expression and pregnancy outcomes (Vafaei et al., 2020). Further experimental studies are needed to explore additional genes and aspects of pregnancy, shedding light on the role of Wi-Fi radiation in fertility and pregnancy.

In our previous reports, we demonstrated that RFR exposure alters mitochondrial function, induces oxidative stress in the testis, causes DNA damage in testicular cells, retards the germ cell transformation rate during spermatogenesis, delays the sperm cell cycle, and lowers sperm count in exposed Swiss albino mice (Pandey and Giri, 2017; 2018). In the present study, we show that RFR exposure to F0 males may not significantly impact reproductive performance or success; however, it adversely affects the sperm-producing capacity of offspring. This effect may be attributed to the involvement of epigenetic factors, potentially leading to permanent defects in mice testis or altered expression of key genes involved in male reproduction. Nevertheless, this aspect is not evaluated in the present study and serves as a potential avenue for future investigations. Our results underscore a pressing issue faced by modern society, marked by a consistent increase in infertility incidences among young adults and high rates of reproductive cancers. Epigenetic modulations play a crucial role in regulating cellular functions and contributing to pathological conditions. Epigenetic alterations may induce permanent changes in gene expression that can be transferred to subsequent generations. Kumar et al. (2021) demonstrated a frequency-dose-dependent increase in DNA and histone methylation, leading to subsequent changes in gene expression in the hippocampus of Wistar rats after 2 hours of exposure for 1 month. Mokarram et al. (2017) reported that RFR exposure modulates the methylation pattern of estrogen receptors in rats, potentially increasing the incidence rate of colorectal cancer. In the present study we propose that RFR exposure led to the change in epigenetic imprinting in male germline that resulted in the permanent effects such as thinning of germinal epithelium and low sperm count in F1 generation male.

## 4. Material & Methods

### 4.1. Animals

Male Swiss albino mice, aged 8–10 weeks and weighing between 22–25 g, were procured from the Pasteur Institute located in Shillong, Meghalaya, India. These mice were maintained under standard laboratory conditions throughout the entire experimental period. Ad libitum access to food pellets (Amrut Laboratory Animal Feeds, New Delhi, India) and water was provided to the mice. The experimental protocols involving animal subjects outlined in this study received approval from the Institutional Ethical Committee of Assam University, situated in Silchar, Assam, India. The approval permit number is IEC/AUS/2013037, dated 20 March 2013. This ensures adherence to ethical standards and guidelines in the conduct of animal experimentation.

### 4.2. Experimental Design and exposure conditions

The animals were divided into two groups, comprising six animals each: CON (control, unexposed) and RFR (exposed for 3 hours twice daily for 35 days). The RFR exposure system utilized a generator-amplifier set-up configured to produce an electromotive force similar to that emitted by global system for mobile communications (GSM) mobile phones, specifically at 902.4 MHz with an average power of 250 mW pulsed at a frequency of 217 Hz. The antenna of the generator was connected via RF amplifier (Mini circuits LZY-2) to a vector signal generator (Rohde and Schwarz SMIQ 06B; Munich, Germany). Throughout the exposure period, animals were allowed to move freely within the cage, which was a chromium-nickel metal container designed for protection from possible external telemetric exposure. The measured power density ranged from 2.717 to 0.284 W/m2, accounting for the minimum and maximum distance of the animal from the antenna (4.5–22.5 cm). The output power from the transmitter was regularly monitored using an RF power meter (Hewlett-Packard 437B; Palo Alto, California, USA). This exposure set-up ensured constant and reliable RF signals throughout the experiments. The specific absorption rate was determined according to the method described by Durney et al. (1984), ranging between 0.0516 and 0.0054 W/kg. At the end of the experimental period, all animals were sacrificed, and tissues of interest were harvested. For mating experiments, F0 males were placed with naïve, unexposed females in different cages in a 1:2 ratio for seven days, followed by a 35-day RFR exposure period. Subsequently, the males were removed from the cages, and pregnant females were observed and maintained until delivery. After delivery, pups were counted and observed daily until 21 days. On the 21st day, male and female pups were segregated, with male mice observed and maintained for six weeks before being sacrificed for endpoint evaluation in adulthood.

### 4.3. Scanning Electron microscopy for sperm defects

The epididymal sperms were preserved in Karnovsky’s fixative at 4°C for 24 hours, followed by two washes with 0.1M Sodium Cacodylate buffer at hourly intervals. Subsequently, the specimens were affixed onto brass stubs using adhesive tape, air-dried at 26°C, and coated with gold vapors. The prepared specimens were then examined using a JEOL (JSM-6360) scanning electron microscope, with electron accelerating voltages ranging between 10-20 keV, as per the methodology outlined by Dey et al. (1989).

### 4.4 Comet Assay

The alkaline comet assay was performed on sperm and bone marrow cells using a modified approach based on the methodology outlined by Singh et al. (1998), incorporating adjustments as described by Tice et al. (2000). Cell suspensions were prepared in 1 ml of ice-cold PBS at pH 7.4. Each sample, at a concentration of 2 × 10^4 cells/ml, was mixed with 1% low melting point agarose (LMPA) in a 1:10 ratio. Subsequently, 85 μl of the mixture was spread on frosted slides pre-coated with 1% normal melting agarose (NMA), followed by an additional layer of 90 μl of 0.5% LMPA and chilling on an ice pack. The slides were maintained at 4 °C for 2 hours in an ice-cold lysis buffer (composed of 2.5 M NaCl, 100 mM Na2EDTA, 10 mM Trizma base, 1% Triton X 100, and 10% DMSO; pH 10). Afterward, the slides were immersed in an electrophoresis buffer (300 mM NaOH, 1 mM Na2EDTA; pH 13.5) for 20 minutes at 4 °C, followed by electrophoresis under constant voltage (24 V; 300 mA) and a temperature of 4 CC. Finally, the slides were neutralized with 0.4 M Tris buffer (pH 7.5) for 5 minutes (repeated three times), stained with ethidium bromide (20 μg/L), and subjected to comet scoring using Komet 5.5 software (Kinetic Imaging, UK) connected to a fluorescence microscope Leica, DM 2000 (Leica, Wetzlar, Germany). For each animal, 50 cells per slide were analyzed. The damage index (DI) and damage frequency were calculated by categorizing cells into five classes (0, 1, 2, 3, and 4) with corresponding tail lengths of 0, >0–5, >5–10, >10–20, and >20 µm, respectively, as described previously (Kahl et al., 2012). Formula used for DI (mean):

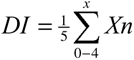

where ‘X’ is the class number and ‘n’ is the percentage of cells in each class.

### 4.5. Sperm Viability test by Flow cytometry

The investigation of live and dead sperm cell populations was conducted using cells obtained from the cauda epididymis of mice, employing propidium iodide (PI) dye. PI, with its ability to intercalate between the double-stranded DNA of deceased cells, facilitated the discrimination between viable and non-viable cells. The cauda epididymis per mouse was minced in 1 ml of PBS containing 0.3% NP-40, gently releasing sperm cells into the buffer. Subsequently, the samples were filtered through a nylon filter and centrifuged at 400 xg for 5 minutes at room temperature.The cells were then permeabilized and stained with PI/RNAse staining buffer (0.5 ml per sample) from BD Biosciences, US. Following an incubation period of 15 minutes at room temperature in the dark, the samples were vortexed and analyzed on a BD Accuri C6 flow cytometer (BD Biosciences, US), recording 50,000 events per sample. Two distinct populations of cells were identified in the cytogram, representing the live and dead sperm populations, respectively.

### 4.6. Mating Experiment

Swiss albino male mice were partitioned into two groups: a control group that remained unexposed (10 males) and an RFR-exposed group (10 males) subjected to a daily exposure of 6 hours for 35 consecutive days. Following the exposure period, each RFR-exposed male was placed in a cage with two unexposed females for a cohabitation duration of 7 days (1 male and 2 females per cage). Following this cohabitation period, a segregation phase of 21 days ensued, during which males were removed from the cages, and females were monitored for pregnancies. Throughout the segregation period, data pertaining to pregnancy outcomes were recorded at various time points. This included the counting of litters, and the maintenance of live pups until weaning. The data collected from different time points allowed for a comprehensive analysis of pregnancy outcomes in response to the experimental conditions.Data was recorded at different time points for analysing pregnancy outcomes. Following parameters were calculated for control and RFR exposed group as described :

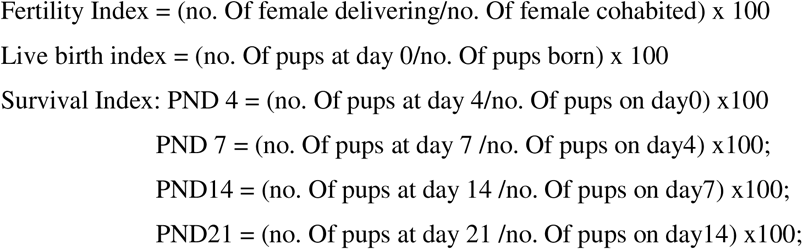

Also, number of pups per litter and sex ratio were recorded on 21PND in F1 generation offsprings.

### 4.7 Sperm count and sperm defects in F1-male

The cauda epididymides were meticulously isolated and finely minced with scissors in 1 mL of 0.9% sodium chloride (NaCl), and the resulting suspension was filtered through a nylon mesh sieve. The filtrate was then diluted at a 1:20 ratio. Subsequently, the number of sperm in five diagonal squares of a Neuberger hemocytometer was counted, and the measured sperm count was expressed as the total number per milliliter (10^6), taking into account chamber and dilution factors.

Simultaneously, utilizing the same epididymal filtrate, a morphological assessment of sperm was conducted for each animal. A drop of filtrate was placed on each slide, immediately smeared, and fixed with methanol. The slides were subsequently stained with 2% Eosin Y stain to facilitate the identification of morphological distinctions between normal and altered sperm. A total of one thousand sperm per animal were scored according to the criteria outlined by Wyrobek and Bruce (1978).

### 4.8 Testis Histology of F1-male

For each animal, one testis was fixed in 10% buffered formalin, followed by dehydration and embedding in paraffin. Subsequently, 5 mm tissue sections were obtained from the paraffin blocks and stained with hematoxylin and eosin (Lillie, 1965). The prepared slides were then examined using a light microscope, specifically the Leica DMLS model (Leica, Wetzlar, Germany). The percentage of atrophied seminiferous tubules per mouse was calculated based on the examination of the tissue sections. Additionally, the percentage of epithelium area per seminiferous tubule was determined using Image J1 image analyzer software (National Institutes of Health, Bethesda, Maryland, USA). A minimum of 30 seminiferous tubules per testis were subjected to analysis. This approach facilitated a comprehensive evaluation of the testicular morphology and provided quantitative data on atrophy and epithelial changes in the seminiferous tubules.

### 4.9 Statistical Analysis

Statistical analysis involved the utilization of one-way ANOVA to compare changes in means across the experimental groups. Subsequently, Tukey’s post hoc analysis was applied to ascertain the significance of differences in means between specific experimental groups. Student’s T-test was used to analyse the comet data between two experimental groups. The statistical analyses were conducted using GraphPad Prism 4 software, with significance determined at a 95% confidence interval (CI) level. Variances were considered significant at a ‘P’ value less than 0.05, indicating a threshold for statistical significance in the observed results.

## 5. Conclusion

This study represents a pioneering effort, being the first to elucidate the involvement of epigenetic factors in the Radiofrequency Radiation (RFR) exposure-induced reduction in sperm count in offspring through the male germline. Exposure of male Swiss albino mice to RFR at a frequency of 900 MHz induces oxidative stress, leading to DNA damage and abnormalities in sperm cells. Subsequently, when these exposed male mice are mated with unexposed, naïve females, the F1 generation mice exhibit low sperm count and depletion of the germinal epithelium in adulthood. These findings underscore the significance of altered gene expression in the testis of F1 males, potentially resulting in permanent changes such as reduced sperm count and thinning of the germinal epithelium in the testis of F1 male mice.

## Acknowledgement

Authors are grateful to the UGC for Phd fellowship to NP. The infrastructural and research facility provided by UGC-SAP supported Department of Life Science and Bioinformatics is acknowledged. Authors are thankful to SAIF-NAHU and Dr Arun Yadawa (Department of Zoology, NEHU, Shillong) for providing Scanning electron microscopy facility and histology facility respectively.

## Ethics Statement

All the authors approve the manuscript

